# A characterization of cardiac-induced noise in R_2_* maps of the brain

**DOI:** 10.1101/2022.12.19.521028

**Authors:** Quentin Raynaud, Giulia Di Domenicantonio, Jérôme Yerly, Thomas Dardano, Ruud B. van Heeswijk, Antoine Lutti

**Author notes:** **Correspondence** Antoine Lutti, Laboratory for Research in Neuroimaging, Department for Clinical Neuroscience, Lausanne University Hospital, Ch. de Mont-Paisible 16, CH-1011 Lausanne.

## Abstract

**Purpose:** Cardiac pulsation increases the noise level in brain maps of the transverse relaxation rate R_2_*. Cardiac-induced noise is challenging to mitigate during the acquisition of R_2_* mapping data because its characteristics are unknown. In this work, we therefore aim to characterize cardiac-induced noise in brain maps of the MRI parameter R_2_*.

**Methods:** We designed a sampling strategy to acquire multi-echo 3D data in 12 intervals of the cardiac cycle, monitored with a fingertip pulse-oximeter. We measured the amplitude of cardiac-induced noise in this data and assessed the effect of cardiac pulsation on R_2_* maps computed across echoes. The area of k-space that contains most of the cardiac-induced noise in R_2_* maps was then identified. Based on these characteristics, we introduced a tentative sampling strategy that aims to mitigate cardiac-induced noise in R_2_* maps of the brain.

**Results:** In inferior brain regions, cardiac pulsation accounts for R_2_* variations of up to 3s^-1^ across the cardiac cycle, i.e. ∼35% of the overall variability. Cardiac-induced fluctuations occur throughout the cardiac cycle, with a reduced intensity during the first quarter of the cycle. 50-60% of the overall cardiac-induced noise is localized near the k-space centre (k < 0.074 mm^-1^). The tentative cardiac noise mitigation strategy reduced the variability of R_2_* maps across repetitions by 11% in the brainstem and 6% across the whole brain.

**Conclusion:** We provide a characterisation of cardiac-induced noise in brain R_2_* maps that can be used as a basis for the design of mitigation strategies during data acquisition.

## 1. Introduction

MRI relaxometry consists of estimating the value of the MRI parameters that drive signal intensities in MR images^1–3^. Relaxometry data exhibit a lower dependence on acquisition parameters and scanner hardware than conventional structural MRI data, leading to increased reproducibility in multi-centre studies^4,5^. Maps of the transverse relaxation rate (R_2_* =1/T_2_^*^) are computed from gradient-echo MR images acquired at multiple echo times. R_2_* relaxation is driven by spin-spin interactions and microscopic magnetic field inhomogeneities that arise from magnetic material within the tissue^6,7^. Therefore, R_2_* correlates with e.g. iron and myelin concentration in brain tissue^8–11^, and enables the monitoring of disease evolution in Parkinson’s disease^12,13^, multiple sclerosis^14^, and Alzheimer’s disease^15^.

Cardiac pulsation leads to instabilities in brain MRI data that reduce the sensitivity of R_2_* estimates to brain disease in neuroscience studies. Cardiac pulsation gives rise to a systolic pressure wave that reaches the brain ∼50ms after onset of the R-wave in an electrocardiogram^16,17^, and 30-130ms prior to the detection of the systolic peak measured with a pulse-oximeter attached to the finger^18–20^. This wave results in pulsatile brain motion due to the expansion of blood vessels^21^, variations in blood flow velocity^17,22–24^, brain tissue deformation^25–28^, CSF motion^29–31^, bulk head motion^32^ and changes in O_2_/CO_2_ concentrations^33,34^. The brain areas primarily affected by cardiac pulsation are inferior brain regions close to large vessels such as the brainstem^35^, cerebellum^36^ and orbitofrontal cortex^31,37^, highly vascularized grey matter regions^31^ and brain regions near the ventricles^37,38^. The effects of cardiac pulsation on MR images arise from the interaction of spin displacements with the amplitude and direction of the gradients of magnetic fields used for image encoding^39^. While laminar flow leads to a net phase shift in MRI data, turbulent or anisotropic flow across an image voxel leads to a distribution of spin phase that results in a net loss in signal amplitude^35^. Cardiac pulsation similarly reduces the BOLD sensitivity of functional MRI data^31,34,35,40^ and leads to bias in measures of the apparent diffusion coefficient^27^.

R_2_* relaxometry requires multi-echo images acquired using oscillating, high-amplitude readout gradients. Because the coupling between encoding gradients and cardiac pulsation accumulates over time^39^, cardiac-induced noise is expected to increase with the echo time, leading to exponential-like effects and thus bias of the R_2_* estimates. Furthermore, because data acquisition takes place over several minutes, raw k-space data points may show variable levels of cardiac-induced noise, leading to aliasing artifacts in the reconstructed images^41,42^. Few data acquisition strategies currently exist that mitigate cardiac-induced noise in R_2_* maps of the brain. On the model of diffusion acquisitions^18,19^, such strategies might consist of adjusting k-space sampling in real-time according to patients’ cardiac pulsation, or of averaging data across multiple samples in the sensitive areas of k-space. These strategies all hinge on a detailed description of cardiac-induced noise in brain R_2_* maps. Such a description includes an assessment of its amplitude, spatial extent in image space and k-space, and timing relative to the cardiac cycle and is not currently available.

Here we provide a complete assessment of cardiac-induced noise in R_2_* relaxometry data of the brain. Using a dedicated and optimized sampling strategy, we acquire multi-echo data in multiple intervals of the cardiac cycle to resolve the effect of cardiac pulsation on the MR signal. From this data, we measure the amplitude of cardiac-induced noise across the cardiac cycle, at all echo times. We assess the effect of cardiac pulsation on estimates of R_2_* computed from the decay of the MRI signal across echoes. Subsequently, we identify the area of k-space that contains most of the cardiac-induced noise in R_2_* maps. From these results, we introduce a tentative sampling strategy that aims to mitigate cardiac-induced noise in R_2_* maps of the brain.

## 2. Methods

Data acquisition was performed with a 3T MRI scanner (Magnetom Prisma, Siemens Healthcare, Erlangen, Germany) equipped with a 64-channel head-neck coil. The study was approved by the local Ethics Committee and all participants gave their written informed consent prior to participation.

The acquisition protocols are presented in Table 1. All MRI acquisition protocols included an MP-RAGE^43^ image for segmentation and anatomical reference (1mm^3^ isometric resolution, TR/TE = 2000/2.39ms, GRAPPA^44^ acceleration factor 2 with 24 reference lines, RF excitation angle = 9°, acquisition time 4:16 minutes).

**Table 1:** MRI acquisition protocols. CG=cardiac gating

| Experiment | Data type | Pulse sequence | #echoes | Spatial Resolution [mm<sup>3</sup>] | Acquisition time [min] |
| --- | --- | --- | --- | --- | --- |
| Characterization of $R_2^*$ changes across the cardiac cycle | 5-dimensional (x-y-z-echoes-cardiac) | FLASH<br>Optimized k-space sampling | 15 | 2x4x4 | 56:21 |
|  | Coil Sensitivity | FLASH<br>Linear cartesian k-space sampling | 1 | 4x4x4 | 0:16 |
|  | T1-weighted image | MPRAGE | 1 | 1x1x1 | 4:16 |
| Informed cardiac-induced noise mitigation strategy | $R_2^*$ mapping (CG on/off) | FLASH<br>Linear cartesian k-space sampling | 15 | 1x2x2 | 7.34 (CG off)<br>8:48±0:04 (CG on) |
|  | T1-weighted image | MPRAGE | 1 | 1x1x1 | 4:16 |

### 2.1. Characterization of R_2_* changes across the cardiac cycle

#### 2.1.1. MRI protocol

Continuous low-resolution multi-echo data was acquired using a custom-written 3D fast low-angle shot (FLASH) sequence on five adult participants (2 females, 33±7 years old) to characterize R_2_* variability across the cardiac cycle. 15 echo images were acquired with a bipolar readout^4^ (repetition time TR=40ms; echo times TE=2.34ms to 35.10ms with 2.34ms spacing, RF excitation flip angle = 16°. The readout direction was set along the head-feet direction. Image resolution was 2mm along the readout direction and 4mm along the phase-encode directions - sufficient to fully capture cardiac-induced fluctuations, because physiological noise scales with signal amplitude and is strongly reduced in high k-space frequency regions required to achieve high spatial resolution^45^.

Our strategy for the acquisition of 3D multi-echo data across the cardiac cycle was inspired by recent developments in high-dimensional heart and brain imaging^22,46–49^, where the cardiac cycle is resolved by pooling k-space data from multiple heartbeats into separate images for each phase of the cardiac cycle. Data were acquired continuously while the cardiac rhythm of the participants was being recorded using a pulse-oximeter attached to the finger. Data acquisition was thus not synchronized with the heart rates of the participants and was conducted using a sampling scheme that was optimized to provide samples at different phases of the cardiac cycle for each k-space location, and to simultaneously mitigate non-cardiac spurious effects such as head-motion, breathing or swallowing (strategy 2 in Supplementary Material). In brief, this scheme consisted of a linear Cartesian sampling within a predefined kernel, i.e. a subset of k-space along the two phase encoding directions. The kernel size was set to 30 by 2 along the fast and slow phase-encoding directions, respectively. 30 samples were consecutively acquired for each phase encoding combination.

The data was binned retrospectively according to the phase of the cardiac cycle at the time of its acquisition, leading to five-dimensional (5D) datasets with three spatial dimensions, one echo-time dimension and one cardiac phase dimension (Figure 1A). We divided the cardiac cycle into 12 bins as a trade-off between sufficient resolution of the systolic period (∼300ms duration^16,31^) and sufficient k-space data in each bin for routine image reconstruction. If multiple samples of the same k-space point were present in a cardiac phase bin, they were averaged. The cardiac phase was set to zero at the peak of the pulse-oximeter signal.

**Figure 1:**
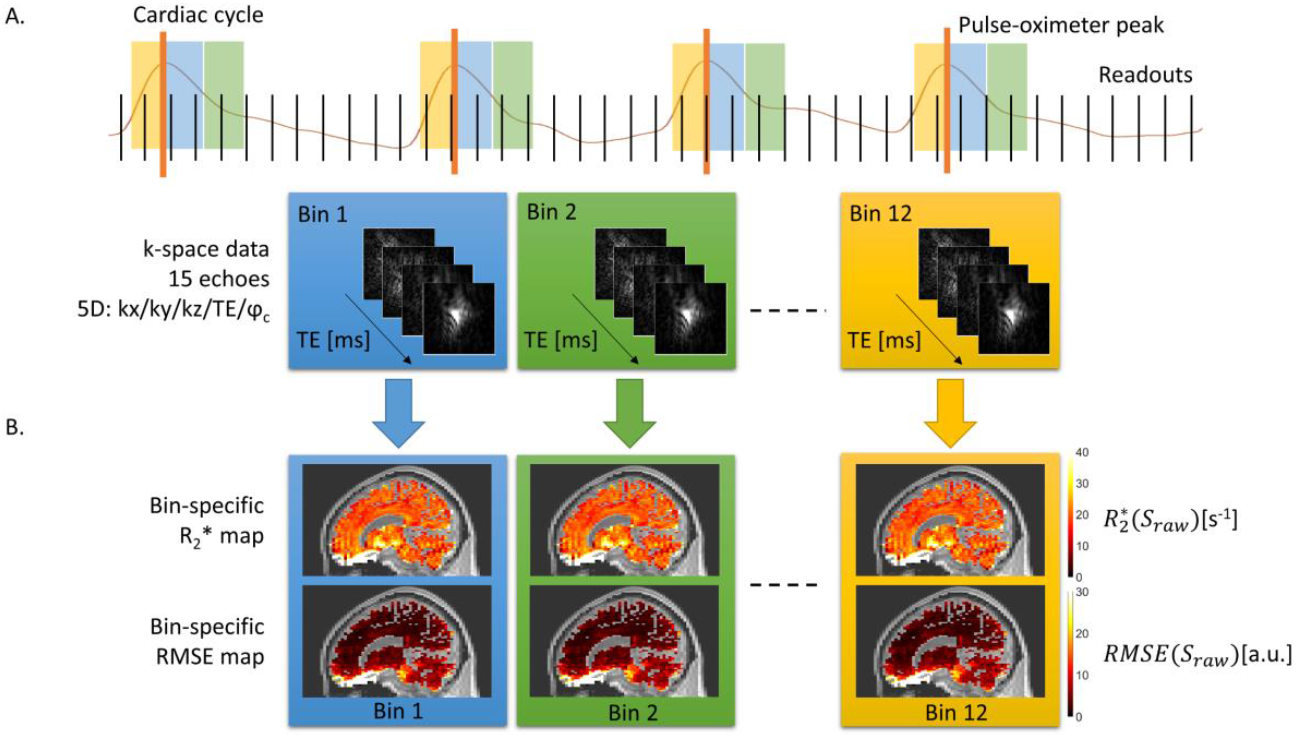
Schematic representation of the acquisition of 5D datasets for the characterization of cardiac-induced noise in brain R_2_* maps. (A.) Multi-echo data was acquired continuously for 1 hour while the cardiac rhythm of the participants was recorded using a pulse-oximeter attached to the finger. The data was binned retrospectively according to the phase of the cardiac cycle at the time of its acquisition, leading to k-space datasets with three spatial dimensions (readout and two phase encoding directions), one echo-time dimension and one cardiac phase dimension. (B.) R_2_* maps were computed after image reconstruction, from the regression of the log signal with the corresponding echo times. The noise level of the R_2_* estimates was calculated as the root-mean-squared error (RMSE) between the MR signal and the R_2_* fit.

To compute the coil sensitivity profiles used for image reconstruction^50^, two 3D FLASH datasets were acquired in each participant with a 4×4×4mm^3^ spatial resolution, with both head and body coils for signal reception (TR/TE=5.72ms/2.34ms, excitation flip angle=6°, acquisition time 16s)^51^.

#### 2.1.2. Data processing and image reconstruction

Following data acquisition, the <10% of missing k-space points in the 5-dimensional datasets were reconstructed using SPIRiT^52^ (http://people.eecs.berkeley.edu/~mlustig/Software.html). The data was inverse Fourier transformed along the fully sampled readout direction and each 2D slice was reconstructed separately for each echo image and cardiac phase. To optimize the accuracy of the reconstruction, the 2D SPIRiT kernel was calibrated using all the k-space data for each bin of the cardiac cycle, after linear interpolation of the missing k-space data from the neighbouring bins. The kernel size was [3x3] and 30 iterations were performed. After SPIRiT reconstruction, the head-foot direction was Fourier-transformed back into k-space data.

Brain images were reconstructed from the completed 5-dimensional k-space datasets using SENSE^50^ with an acceleration factor of 1. These images contained signal from areas outside and below the brain (e.g. tongue, mouth, neck) along the readout encoding direction (orientation: head-feet). Cardiac-induced noise from these areas might alias in the 2D plane of the two phase encoding directions, but cannot alias into the brain along the readout direction due to the high readout bandwidth. Therefore, these noise contributions are of no interest for the characterization of cardiac-induced noise in brain images and the 5D images were trimmed below the medulla along the readout direction.

#### 2.1.3. Modelling of cardiac-induced k-space fluctuations

Consistent with common models of physiological noise^40,45,53^, cardiac-induced fluctuations of the complex MRI signal across the cardiac cycle were modelled at each k-space location and each echo separately using 2^nd^-order Fourier series of sinusoidal basis functions (Figure 2A):

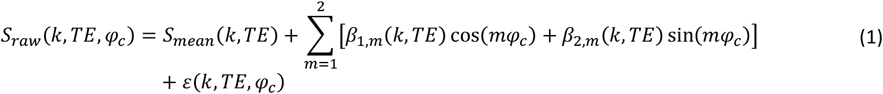

where *S*_*raw*_ is the raw 5D MR data, *S*_*mean*_ is the mean signal across the cardiac cycle, *φ*_*c*_ is the phase of the cardiac cycle, *TE* is the echo time of the data and ε is the residual. β_1,*m*_ and β_2,*m*_, the complex weights of the *m*^th^-order components, were estimated from the real and imaginary parts of *S*_*raw*_ instead of its phase and magnitude to improve resilience against low signal-to-noise^45^. From the estimates of *β*_1,*m*_ and *β*_2,*m*_, the modelled cardiac-induced fluctuations of the MRI signal *S*_*cardiac*_ were computed as follows:

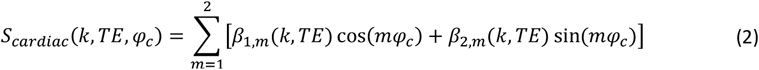

**Figure 2:**
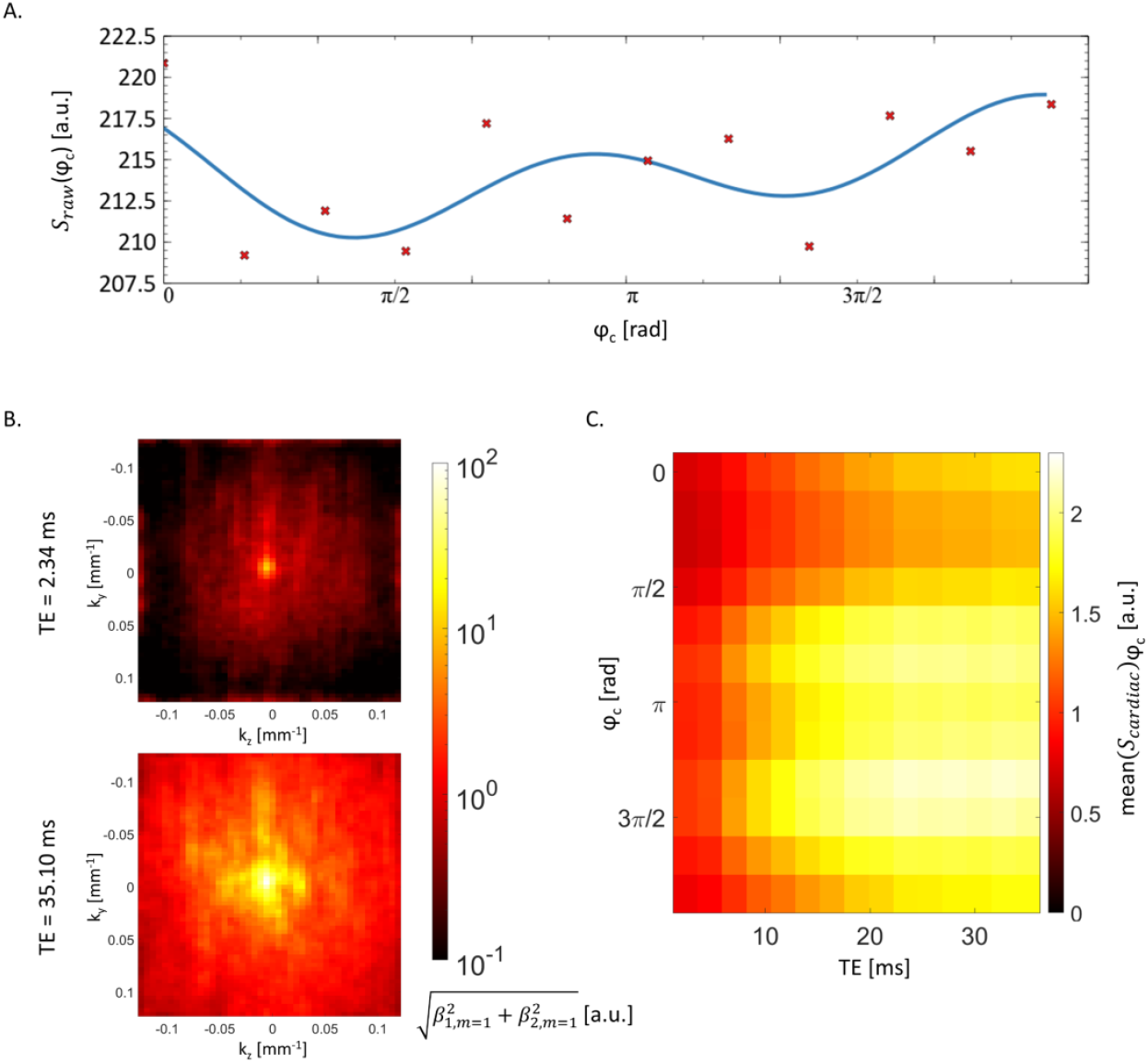
Cardiac-induced noise in the multi-echo data. (A) Example fluctuation of the raw signal S_raw_ across the cardiac cycle at one k-space location of the last echo data (TE = 35.10 ms) (red dots). The blue solid line shows the corresponding modelled cardiac-induced noise S_cardiac_ + S_mean_ across the cardiac cycle. (B) K-space distribution of the amplitude of the 1^st^ Fourier component of the modelled cardiac-induced fluctuations, averaged along the readout direction and participants. (C.) Average deviation of S_cardiac_ from its mean value across the cardiac cycle, averaged across k-space and participants.

Similarly, the non-cardiac fluctuations of the raw MR signal *S*_*no cardiac*_ were computed as follows:

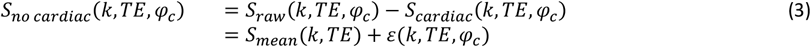

#### 2.1.4. Characterization of R_2_* changes across the cardiac cycle

To analyse the effects of all noise sources on R_2_* estimates, we computed R_2_* maps for each cardiac bin from *S*_*raw*_. We applied a Fourier transform along the three k-space dimensions of *S*_*raw*_. For each phase of the cardiac cycle, R_2_* maps were computed from the resulting images from a regression of the log signal with the corresponding echo times^54^ 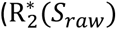, figure 1B). The noise level on the R_2_* estimates was calculated as the root-mean-squared error (RMSE) between the MR signal and the R_2_* fit (*RMSE*(*S*_*raw*_)).

This procedure was also performed on *S*_*cardiac*_+ *S*_*mean*_ to analyse the effects of the modelled cardiac pulsation, leading to estimates 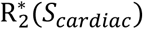 and RMSE(*S*_*cardiac*_). To analyse the changes in R_2_* across the cardiac cycle due to non-cardiac noise sources, this procedure was conducted on *S*_*no cardiac*_(*k,TE, φ*_*c*_), leading to estimates 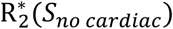 and *RMSE*(*S*_*no cardiac*_).

#### 2.1.5. Determination of the k-space region sensitive to cardiac-induced fluctuations

To characterize the distribution of cardiac-induced noise across k-space, we removed the modelled cardiac-induced noise from sub-regions of the 5D raw data and measured the impact on the standard deviation (SD) of R_2_* across the cardiac cycle. We computed the hybrid k-space matrix:

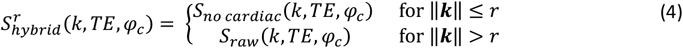

where ‖*k*‖ is the distance to the center of a given k-space location along the two phase encoding directions, and *r* a given radius. 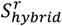 contains no cardiac-induced noise inside a circular area of radius *r*, and noise of all sources outside. For values of *r* from 0% to 100% of k-space extent with step size 1%, R_2_* maps were computed across the cardiac cycle using the procedure outlined in 2.1.4. The changes in R_2_* across the cardiac cycle were used to measure the effect of cardiac-induced noise in an area outside *r*, and were compared to the effect of all sources of noise present in the data.

The same procedure was repeated using the hybrid k-spaces that only contained cardiac noise, so that the contributions of cardiac and non-cardiac fluctuations could be compared:

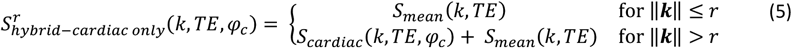

The changes in R_2_* across the cardiac cycle allowed the estimation of the effect of cardiac-induced noise alone in an area of k-space outside *r*.

### 2.2. Informed cardiac-induced noise mitigation strategy

Once the sensitive k-space regions were determined as described above, we implemented a mitigation strategy similar to cardiac gating that is widely used in diffusion MRI^18–20^ that involves suspending data acquisition during detrimental periods of the cardiac cycle.

Data were acquired on seven adult participants (6 females, 32±7 years old) using a multi-echo 3D FLASH sequence with a linear Cartesian trajectory (repetition time TR=40ms; echo times TE=2.34ms to 35.10ms with 2.34ms spacing, RF excitation flip angle = 16°). The voxel size was 1×2×2mm^3^, similar to brain R_2_* maps acquired in clinical protocols^55–57^.

With our implementation, the time window for data acquisition was limited to the first quarter of the cardiac cycle that followed detection of the R-wave with the pulse-oximeter, because this period was observed to contain the least cardiac-induced noise (see Results section). To minimize the increase in scan time due to the suspension of data acquisition, cardiac gating was only effective for the acquisition of the subset of k-space that contains most of the cardiac-induced noise (see Results section). This corresponds to 22% of k-space at the resolution of the 5D acquisition (2x4x4mm^3^), but only 5.5% of k-space with the resolution used here (1x2x2mm^3^). Note that this implementation of cardiac gating for 3D FLASH sequences maintained RF excitation during periods of suspension to preserve the steady state of the magnetization^58^. The acquisition time was 8:48±0:04 and 7:34 minutes with and without cardiac gating respectively. Data acquisition was conducted 3 times for both conditions in a randomized order.

The effect of cardiac gating on brain R_2_* maps was measured from the repeatability of the maps across 3 repetitions and from the maps of RMSE, the goodness-of-fit of the MRI signal with the R_2_* fit.

### 2.3. Image analysis

Image reconstruction and data analyses were performed using bespoke analysis scripts written in MATLAB (version 2017a, The MathWorks, Natick, MA). The impact of cardiac-induced noise was assessed in four different regions of interest (ROIs): brainstem, cerebellum, whole brain, and noisy non-brain voxels. The noisy non-brain region was designed to include areas outside brain tissue with high levels of cardiac-induced noise (e.g. blood vessels, CSF). With standard acquisition strategies, this noise might alias across images and enhance the effective noise level in brain voxels. As the acquired 3D FLASH data did not allow for accurate delineation of blood vessels, we devised an ad-hoc procedure to compute a mask composed of voxels outside brain tissue that exhibit a high level of cardiac-induced noise. The voxels within this mask showed a combined grey and white matter probability below 0.1, and variations of the modelled cardiac-induced noise *S*_*cardiac*_ across the cardiac cycle in the last echo image above the average value within the brain.

Image coregistration and segmentation were conducted using Statistical Parametric Mapping (SPM12, Wellcome Centre for Human Neuroimaging, London, UK). The MPRAGE images were segmented into maps of grey and white matter probabilities using Unified Segmentation^59^. Whole-brain masks were computed from the grey and white matter segments and included voxels with a combined probability of 0.9 or above. As described in Lutti et al.^60^, regional masks were computed from the grey matter maximum probability labels computed in the ‘MICCAI 2012 Grand Challenge and Workshop on Multi-Atlas Labeling’ (https://masi.vuse.vanderbilt.edu/workshop2012/index.php/Challenge_Details), using MRI scans from the OASIS project (http://www.oasis-brains.org/) and labelled data provided by Neuromorphometrics, Inc. (http://neuromorphometrics.com/) under academic subscription.

## 3. Results

Figure 2A shows an example of fluctuation of the raw MR data *S*_*raw*_ across the cardiac cycle and of modelled cardiac-induced fluctuation *S*_*cardiac*_. The amplitude of *S*_*cardiac*_ is two orders of magnitude larger at the centre of k-space than at the edges (Figure 2B). The average amplitude of *S*_*cardiac*_ increases by a factor 2-2.5 with echo time and varies by a factor 1.5 across the cardiac cycle (Figure 2C). Consistent with previous observations^16–20,61^, the maximum cardiac-induced noise level occurs upon arrival of the systole in the brain (cardiac phase ≈ 3π/4). The first quarter of the cardiac cycle shows a diastolic and reduced level of cardiac-induced noise.

Cardiac-induced noise leads to a variability of 0.59s^-1^ in R_2_* across the cardiac cycle, averaged across the brain (Table 2). In the brainstem and cerebellum, this variability is 0.95s^-1^ and 0.77s^-1^ respectively. Accounting for all sources of noise in the raw MRI data (i.e. 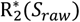), changes in R_2_* of up to 3s^-1^ occur across the cardiac cycle (Figure 3A as well as Supplemental Animation 1 and 2). The modelled changes in R_2_* across the cardiac cycle reflect exponential-like effects of cardiac pulsation on the MRI signal across echoes (Figure 3B). Before removal of cardiac-induced fluctuations (i.e. 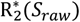), the highest levels of R_2_* variation across the cardiac cycle are observed in inferior brain regions such as the brainstem and cerebellum (Figure 3C and table 2). After removal of cardiac-induced fluctuations (i.e. 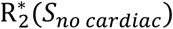), the spatial uniformity of the R_2_* variations is improved. At the regional level, cardiac pulsation accounts for ∼35% (p<0.05) of the overall SD of R_2_* across the cardiac cycle in the brain and 44% (p<0.05) in non-brain voxels (e.g. blood vessels, CSF), respectively 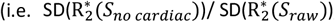, see Figure 3D and Table 2).

**Table 2:** Standard deviation of R_2_* across the cardiac cycle, calculated from the raw MR signal (*S*_*raw*_), the modelled cardiac-induced noise S_cardiac_), the raw MR signal after removal of the modelled cardiac-induced noise (*S*_*no cardiac*_) and two hybrid 5D k-space 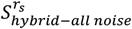 and 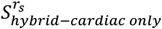, computed from *S*_*raw*_ and *S*_*cardiac*_, by removal of cardiac-induced noise within a circular region of radius *r*_*s*_ = 0 0.074 mm^−1^.

|  | Brainstem | Cerebellum | Whole-brain | Noisy non-brain voxels |
| --- | --- | --- | --- | --- |
| $SD(R_2^*(S_{raw}))_{\varphi_c} [s^{-1}]$ | 1.35±0.29 | 1.04±0.19 | 0.82±0.18 | 3.55±0.53 |
| $SD(R_2^*(S_{cardiac}))_{\varphi_c} [s^{-1}]$ | 0.95±0.09 | 0.77±0.06 | 0.59±0.06 | 2.97±0.23 |
| $SD(R_2^*(S_{no\ cardiac}))_{\varphi_c} [s^{-1}]$ | 0.91±0.24 | 0.65±0.11 | 0.54±0.13 | 2.00±0.31 |
| $SD(R_2^*(S_{hybrid-all\ noise}^{rs}))_{\varphi_c} [s^{-1}]$ | 1.13±0.24 | 0.80±0.12 | 0.66±0.14 | 2.99±0.38 |
| $SD(R_2^*(S_{hybrid-cardiac\ only}^{rs}))_{\varphi_c} [s^{-1}]$ | 0.60±0.09 | 0.43±0.06 | 0.35±0.06 | 2.22±0.23 |

**Figure 3:**
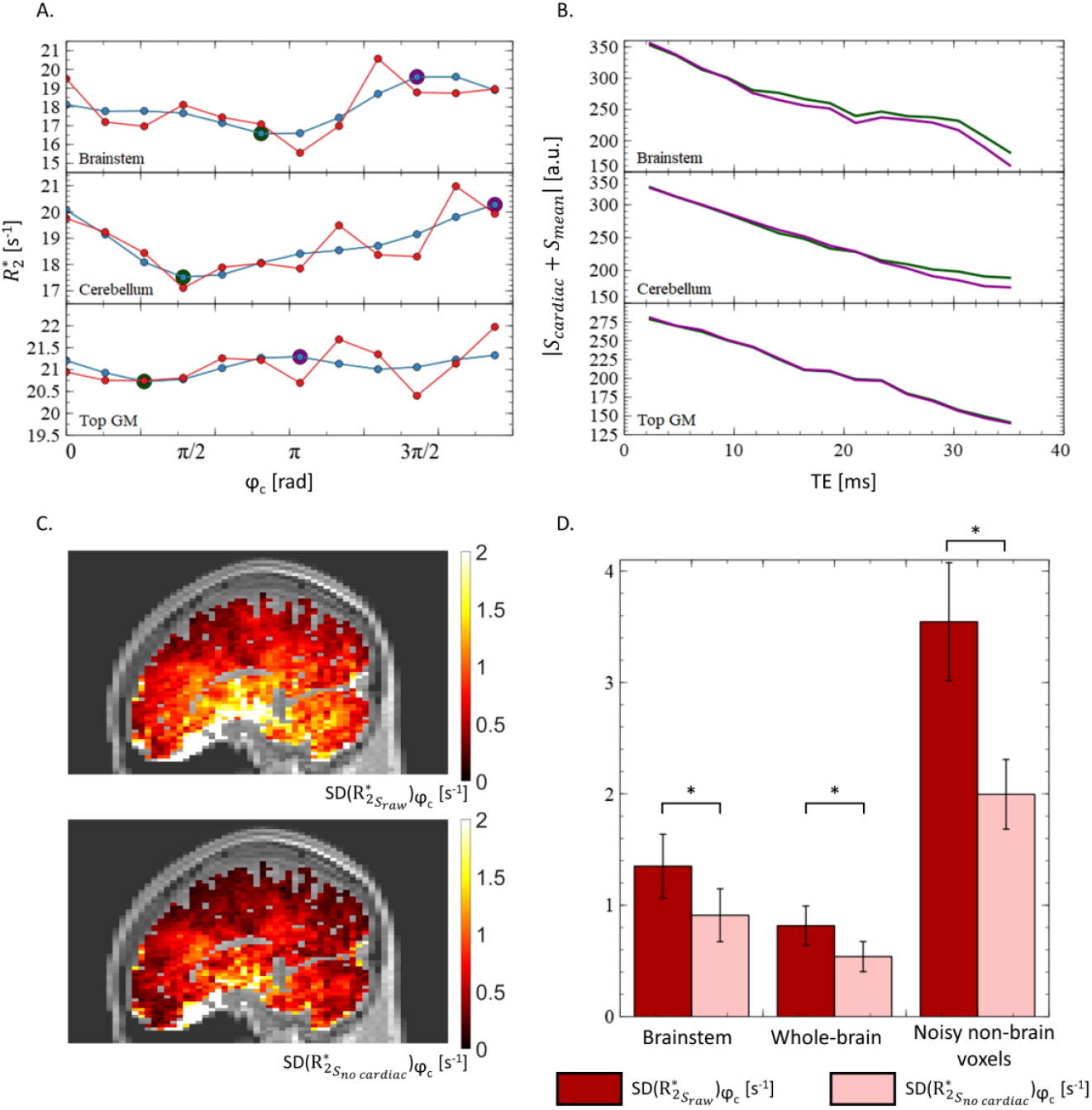
Variability of R_2_* across the cardiac cycle. (A) Example changes in R_2_* across the cardiac cycle due to all noise sources in the raw data 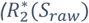, red) and due to the modelled cardiac-induced noise 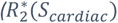, blue). The changes in R_2_* across the cardiac cycle reflect exponential-like effects of cardiac pulsation on the MRI signal. This is illustrated in fig (B) by presenting the magnitude of the modelled cardiac-induced noise S_cardiac_ across echo time for the purple and green points of fig (A), being respectively the maximum and minimum of 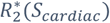 across the cardiac cycle. (C.) Example maps of the variability of R_2_* across the cardiac cycle before and after removal of the modelled cardiac-induced noise 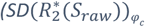 and 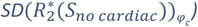 Regional estimates of 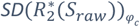 and 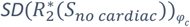 averaged across participants. The difference between those two quantities is significant (p<0.05).

The fitting residuals (RMSE) show variations of up to 40% across the cardiac cycle (Figure 4A). The modelled changes in RMSE across the cardiac cycle reflect non-exponential effects of cardiac pulsation on the MRI signal across echoes (Figure 4B), with a general increase of the RMSE with the echo time. Before removal of the modelled cardiac-induced fluctuations (i.e. *S*_*raw*_), the highest level of variation in RMSE across the cardiac cycle are observed in inferior brain regions such as the brainstem and cerebellum (Figure 4C, see table 3). After removal of cardiac-induced fluctuations (i.e. *S*_*no cardiac*_), the spatial uniformity of the RMSE variations is improved. At the regional level, cardiac pulsation respectively accounts for 29% (p<0.05) of the overall SD of RMSE across the cardiac cycle in the brain and 42% (p<0.05) in non-brain voxels (e.g. blood vessels, CSF) (SD(RMSE(*S*_*no cardiac*_))/ SD(*RMSE*(*S*_*raw*_)), see Figure 4D and Table 3).

**Table 3:** Standard deviation of the fit residuals (RMSE) across the cardiac cycle, calculated from the raw MR signal (S_raw_), the modelled cardiac-induced noise (*S*_*no cardiac*_) and two hybrid 5D k-space 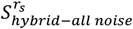 and 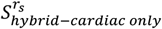, computed from *S*_*raw*_ and *S*_*cardiac*_, by removal of cardiac-induced noise within a circular region of radius *r*_*s*_ = 0 0.074 mm^−1^.

|  | Brainstem | Cerebellum | Whole-brain | Noisy non-brain voxels |
| --- | --- | --- | --- | --- |
| $SD(RMSE(S_{raw}))_{\varphi_c} [\text{a.u.}]$ | 1.52±0.34 | 1.20±0.15 | 0.86±0.16 | 1.85±0.28 |
| $SD(RMSE(S_{cardiac}))_{\varphi_c} [\text{a.u.}]$ | 1.03±0.18 | 0.81±0.15 | 0.58±0.12 | 1.49±0.23 |
| $SD(RMSE(S_{no\ cardiac}))_{\varphi_c} [\text{a.u.}]$ | 1.09±0.29 | 0.85±0.11 | 0.61±0.13 | 1.08±0.16 |
| $SD(RMSE(S_{hybrid-all\ noise}^{rs}))_{\varphi_c} [\text{a.u.}]$ | 1.35±0.29 | 1.03±0.11 | 0.75±0.14 | 1.60±0.22 |
| $SD(RMSE(S_{hybrid-cardiac\ only}^{rs}))_{\varphi_c} [\text{a.u.}]$ | 0.75±0.09 | 0.55±0.07 | 0.41±0.07 | 1.15±0.14 |

**Figure 4:**
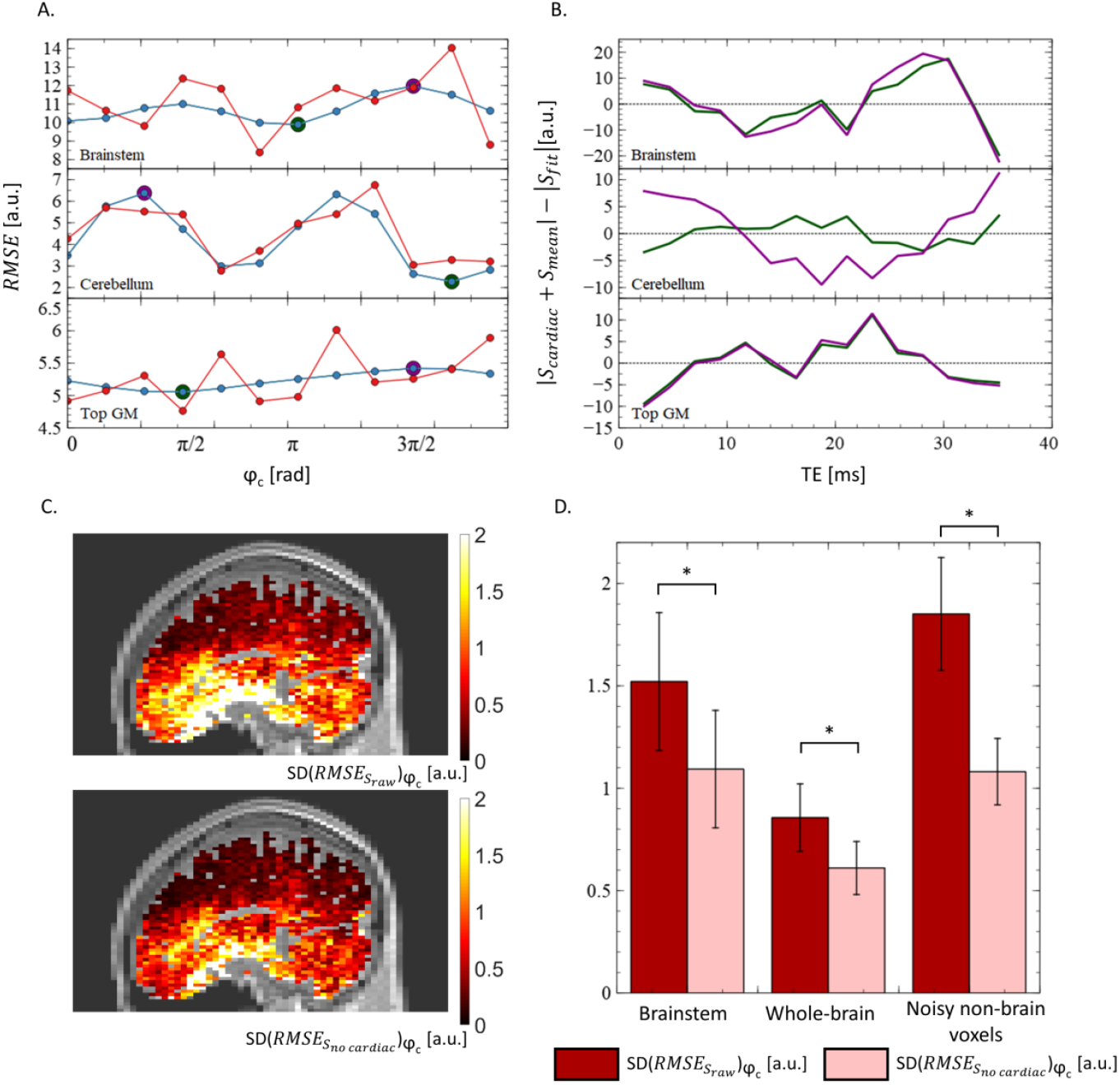
Variability of the R_2_* fitting residuals (RMSE) across the cardiac cycle. (A.) Example changes in RMSE across the cardiac cycle due to all noise sources in the raw data (RMSE(S_raw_), red) and due to the modelled cardiac-induced noise (RMSE(S_cardiac_), blue). The changes in RMSE(S_cardiac_) across the cardiac cycle reflect non-exponential effects of cardiac pulsation on the MRI signal. This is illustrated in fig (B) by presenting the difference between the magnitude of the modelled cardiac-induced noise S_cardiac_ and the magnitude of the exponential fit S_fit_ across echo time for the purple and green points of fig (A), being respectively the maximum and minimum of RMSE(S_cardiac_) across the cardiac cycle. (C.) Example maps of the variability of RMSE across the cardiac cycle before and after removal of the modelled cardiac-induced noise 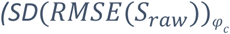 and 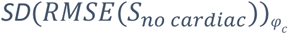 respectively). (D.) Regional estimates 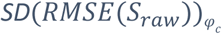 and 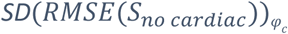 averaged across participants. The difference between those two quantities is significant (p<0.05).

The standard deviation across the cardiac cycle of R_2_* maps computed from 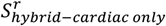 shows a decrease with increasing *r*, with an inflection point near ∼80% of detrended voxels (Figure 5A). From 80 to 100% of detrended voxels, the standard deviation of R_2_* across the cardiac cycle varies as the inverse of the square root of the number of remaining voxels (black curve), indicating that R_2_* variability is dominated by thermal noise^62^. The apparent knee point near the origin delineates the region of k-space where the local variation in the level of cardiac-induced noise is sharp, i.e. the region of k-space sensitive to cardiac-induced noise. Using the Kneedle algorithm^63^, this region was determined to be a circle of radius *r*_*s*_ = 0.074 mm^-1^ that includes ∼22-24% of the central points of k-space (dashed lines in Figure 5A and Figure 5B). Removal of cardiac-induced noise from this region reduces the variability of R_2_* across the cardiac cycle by ∼40% (p<0.05) across the brain, and 25% (p<0.05) in non-brain voxels such as blood vessels or CSF (i.e. 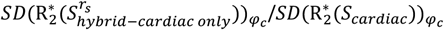). Accounting for all noise sources present in the data (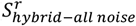), removal of cardiac-induced noise from this region reduces the variability of R_2_* across the cardiac cycle by ∼15-20% (p<0.05) 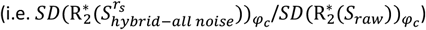, see Table 2 and figure 5C).

**Figure 5:**
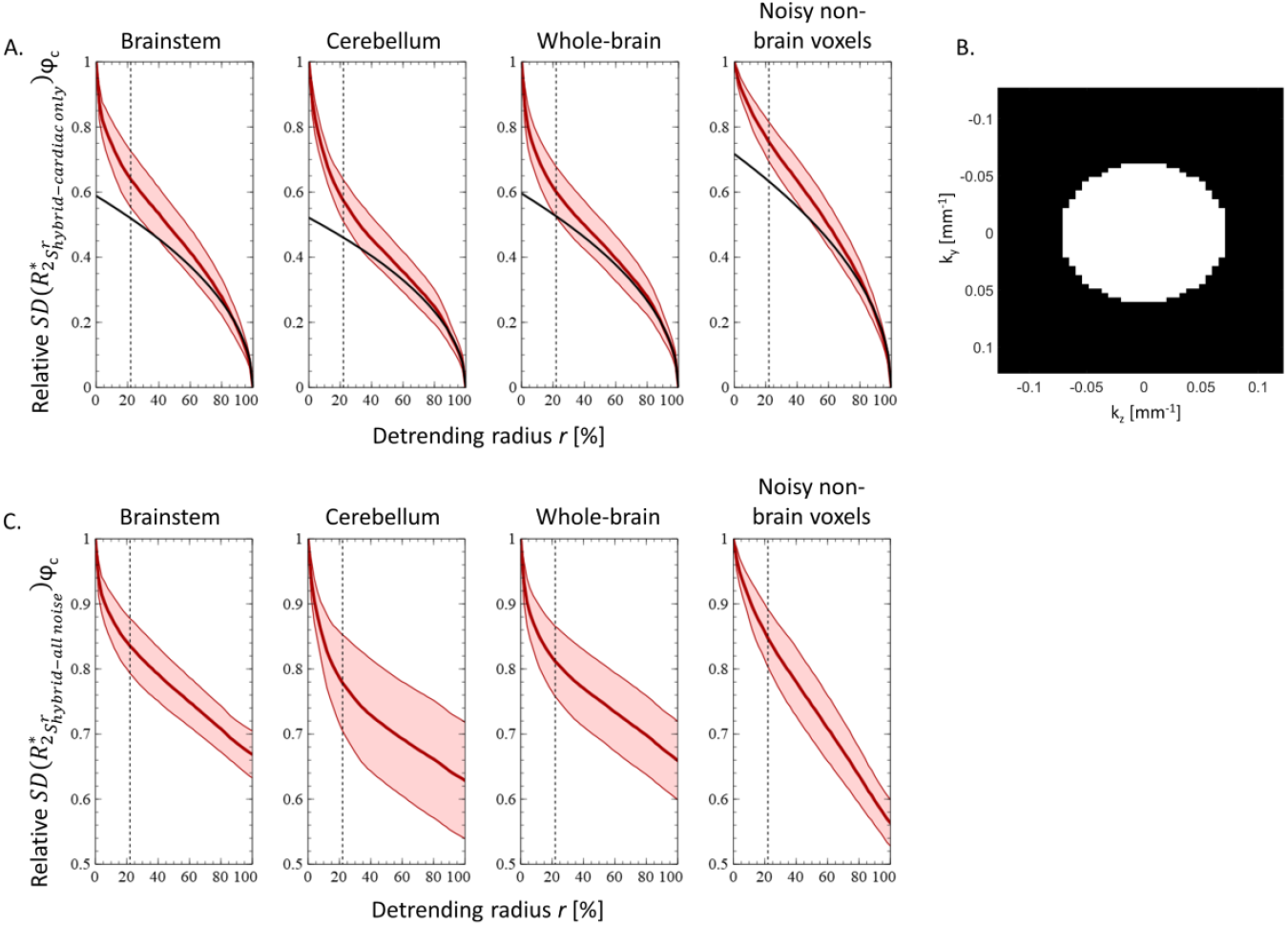
Determination of the k-space regions sensitive to cardiac-induced fluctuations. (A.) Regional estimates of relative R_2_* variability across the cardiac cycle for increasing radius r of 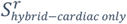. and uniformly distributed noise (black), computed from 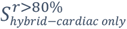. The black dotted line corresponds to the calculated radius of the sensitive region r_s_. (B.) K-space map of the sensitive region covering 22% of k-space. (C.) Regional estimates of relative R_2_* variability across the cardiac cycle for increasing radius r of 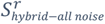.

From the characteristics of cardiac-induced noise presented above, we compared the reproducibility of R_2_* maps obtained with a standard acquisition and with an implementation of cardiac gating (CG) that allowed data acquisition during the first quarter of the cardiac cycle only (Figure 6A). Cardiac gating was only implemented for the acquisition of the sensitive region of k-space, located within a circle of radius *r*_*s*_ = 0.074 mm^-1^. The variability the R_2_* maps across repetitions decreased with the CG sequence, particularly in inferior brain regions: this decrease was 11/8/6% in the brainstem/cerebellum/whole-brain (non-significant, p>0.05) (Figure 6B). Figure 6C shows maps of the RMSE of the fit averaged across repetitions for the standard and CG sequences. Similar to R_2_* variability, the decrease in RMSE with the CG sequence was most pronounced in inferior brain regions: 7% in the brainstem and cerebellum and 3% across the whole brain (non-significant, p>0.05) (Figure 6D). However, aliasing of cardiac-induced noise originating from the circle of Willis along the anterior-posterior phase encode direction can be observed.

**Figure 6:**
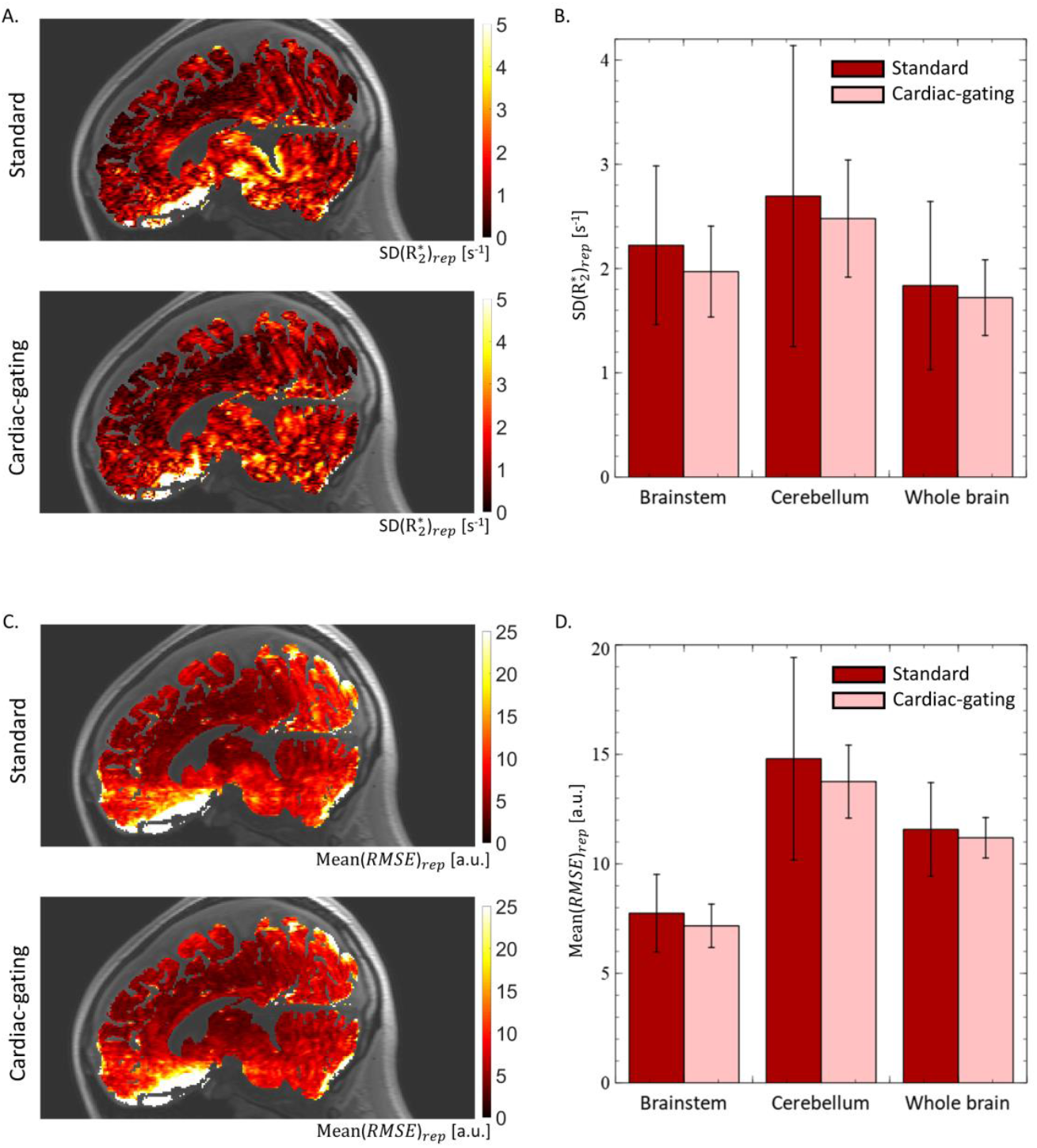
Informed cardiac-induced mitigation strategy. (A.) Example maps of R_2_* variability across repetitions for standard and cardiac-gated sequences. (B.) Regional estimates of R_2_* variability across repetitions, averaged across participants. (C.) Example maps of mean R_2_* fit residuals (RMSE) across repetitions for standard and cardiac-gated sequences (non-significant, p>0.05). (D.) Regional estimates of mean RMSE across repetitions, averaged across participants (non-significant, p>0.05).

## 4. Discussion

We presented an extensive characterization of noise induced by cardiac pulsation in quantitative maps of the MRI parameter R_2_* in the brain. Multi-echo R_2_* mapping data were acquired across the cardiac cycle using a continuous sampling strategy similar to that used for high-dimensional brain and cardiac imaging^22,46–49^. Data acquisition was conducted using a Cartesian sampling kernel optimized to mitigate spurious effects such as breathing- or motion-induced effects and maximize the filling of the multidimensional space of the data. We modelled the effect of cardiac pulsation from the changes of the raw k-space data across the cardiac cycle. We used the modelled cardiac-induced noise to identify the sensitive region of k-space and estimate the effect of cardiac pulsation on the reproducibility of R_2_* estimates. The modelled cardiac-induced fluctuations do not distinguish between the physiological processes that may contribute to the effect, but rather represent an overall measure of cardiac-induced noise in R_2_*-mapping data. From the distribution of cardiac-induced noise in k-space a tentative strategy to mitigate the effect during data acquisition was presented, which leads to an effective reduction in variability of R_2_* maps across repetitions.

The amplitude of cardiac-induced noise increases with the echo time of the data, in line with the expected coupling of motion with the magnetic field gradients used for imaging encoding. This leads to systematic exponential-like effects of cardiac pulsation on the transverse signal decay, and to apparent changes of R_2_* across the cardiac cycle. Variations in R_2_* of 0.59 s^-1^ were observed across the cardiac cycle on average across the brain, reaching 0.95/0.77/2.97s^-1^ in the brainstem/cerebellum/noisy non-brain voxels. Cardiac-induced noise thus accounted for ∼35% of the overall SD of R_2_* across the cardiac cycle in the brain and 44% in non-brain voxels (e.g. blood vessels, CSF). Also, the non-exponential effects of cardiac pulsation on the MRI signal led to an 29% increase of the overall SD of RMSE across the cardiac cycle in the brain and 42% in non-brain voxels.

The amplitude of cardiac-induced noise is strongest near the k-space centre and decreases sharply towards the periphery. The improvements in the reproducibility of R_2_* maps after detrending the modelled cardiac-induced fluctuations over an increasing fraction of k-space allowed us to delineate the region of k-space sensitive to cardiac-induced noise (Figure 5). We found that this region lies within a relatively small radius of 0.074 mm^-1^ from the k-space centre and accounts for ∼40% of the total cardiac-induced fluctuations in the brain and 25% in non-brain voxels (see table 2).

The distribution of cardiac-induced noise in k-space is amenable to the design of mitigation strategies that primarily target the k-space centre. Here, we tentatively chose to mitigate cardiac-induced noise by restricting data acquisition in the target region to the 1^st^ quarter of the cardiac cycle, where the level of cardiac-induced noise is lowest (see figure 2C). This strategy leads to a decrease in R_2_* variability across repetitions of 11%, 8% and 6% in the brainstem, cerebellum, and whole brain, at a cost of 16% scan time increases (Figure 6B). In areas such as the brainstem, this is more efficient than uninformed averaging methods that would require a scan time increase of ∼23% to achieve the same improvements in R_2_* reproducibility. However, it is suboptimal in superior brain regions, where cardiac-induced signal instabilities are not a dominant source of noise. The cardiac-gated acquisitions show improved spatial uniformity. In particular, RMSE maps exhibit a reduced level of aliasing of cardiac-induced noise from the circle of Willis along the anterior-posterior direction.

However, the improvements in the stability of R_2_* maps with the proposed cardiac gating approach only represent a fraction of the changes in R_2_* from the modelled cardiac-induced fluctuations in the 5D data. We are currently investigating alternative strategies that involve the synchronization of data acquisition with cardiac pulsation for Cartesian trajectories, as well as alternative sampling trajectories. Initial results indicate that these strategies provide a much improved mitigation of cardiac-induced noise^66^. The spatial resolution in the assessment of the proposed mitigation strategy was higher than the 5D datasets used for the characterization of cardiac-induced noise, leading to a higher contribution of thermal noise to the overall variability of R_2_* maps. Furthermore, cardiac gating was set to act on a restricted area of k-space that contains ∼40% of the total cardiac-induced noise in brain voxels and ∼25% in non-brain voxels, leaving a large part of cardiac-induced noise in other k-space regions intact. In particular the large remaining cardiac-induced noise in noisy non-brain regions (i.e. blood vessels and CSF) leads to a visible amount of spatial aliasing in the data that propagates into the brain (figure 6A, 6C).

Visual examination of the maps suggests that head motion may have led to image degradation that partly overshadowed the effects of this cardiac noise mitigation strategy. This was confirmed by a quantitative assessment of image quality using a Motion Degradation Index (MDI)^58,60^. Increased MDI values indicative of motion degradation were present in datasets acquired with both standard and cardiac-gated acquisitions (data unshown). It is therefore likely that head motion might have introduced a substantial contribution to the variability of the R_2_* maps between repetitions, leading to a decrease of the relative contribution of cardiac pulsation. The increase in scan time with cardiac gating increases the likelihood of image degradation due to head motion^51,64,65^.

## 5. Conclusion

In this work, we provide a thorough assessment of cardiac-induced noise in brain maps of the R_2_* relaxation rate. Multi-echo data was acquired at regular intervals across the cardiac cycle using an optimized scheme, enabling the analysis of cardiac pulsation effects on brain R_2_* maps. Variations of up to 3/2/1/6s^-1^ in R_2_* were observed across the cardiac cycle in the brainstem, cerebellum, whole-brain and noisy non-brain voxels (e.g. blood vessels, CSF). Cardiac-induced noise accounts for ∼35% of the total R_2_* variability in brain voxels and 44% in noisy non-brain voxels. The amplitude of cardiac-induced noise is strongest near the k-space centre and decreases sharply towards the periphery: the k-space frequencies below 0.074 mm^-1^ (i.e. the centre 22% of k-space at an encoding phase resolution of 4mm) accounts for ∼40% and 25% of the cardiac-induced fluctuations of R_2_* in the brain and in non-brain voxels. This represents ∼20% and 15% of the overall R_2_* variability across the cardiac cycle.

The results of the cardiac-induced noise characterization were used to design a mitigation strategy that reduces efficiently the level of cardiac-induced noise in R_2_* maps of the brain. The proposed cardiac gating strategy suspends data acquisition during detrimental periods of the cardiac cycle. To minimize the increase in scan time that results from the suspension, cardiac gating was only implemented for the acquisition of the subset of k-space that contains most of the cardiac-induced noise. With the proposed strategy, the variability of R_2_* maps was reduced by 11% in the brainstem and 6% across the whole brain, compared to standard acquisition techniques, for an increase in scan time of 13% only.

## Supporting information

animation 1

animation 2

## Data availability statement

The five-dimensional data acquired from one study participant is available online (DOI:10.5281/zenodo.7428605).

The Matlab scripts used for data analysis are available online (DOI:10.5281/zenodo.7446038).

## Acknowledgements

The authors thank Dr. Christopher W. Roy for his help and insights regarding the sampling strategy, as well as all the participants of the study for their time.

## Ethics approval statement

This study was approved by the local Ethics Committee (CER-VD 2016-01284) and all participants gave their written informed consent prior to participation.

## Supplementary material: Optimizing data acquisition for the characterization of cardiac-induced noise

### a. Motivation

A preliminary dataset was acquired in one female participant (31 years old) to check for the presence of non-cardiac spurious effects such as swallowing on the characterization of cardiac-induced noise and to determine the optimal kernel size for the mitigation of these spurious effects. For this dataset, 30 samples were acquired consecutively with linear encoding for each phase encoding combination.

Cardiac-induced noise was modelled using Fourier series decomposition of the signal change across the cardiac cycle. The amplitudes of cardiac-induced noise in k-space across the cardiac cycle show streaks of 1-4 points with high noise amplitude along the fast phase-encode direction, indicating a duration of up to ∼5s during data acquisition (Figure S1). Inspection of the resulting images showed that the source of this effect was primarily located around the mouth and might arise from swallowing.

These observations motivated the design of k-space sampling strategies tailored to mitigate spurious, temporally coherent effects (e.g. swallowing, breathing, head motion, scanner drift) while preserving sensitivity to cardiac-induced noise.

### b. Strategies for the mitigation of spurious effects

The candidate strategies were selected because they distribute spurious effects incoherently across neighbouring k-space locations and were designed around data acquisition within a 2D kernel, i.e. a subset of k-space in the plane of the phase-encode directions. With sampling strategy 1 (Figure S2A), data acquisition is repeated within this kernel before shifting the kernel position by one k-space index along the fast phase-encode direction. With sampling strategy 2 (Figure S2B), the shift of the kernel position takes place after each kernel acquisition and the process is repeated after completing the traversal of k-space along the fast phase-encode direction. Both strategies lead to a total of 30 samples at each k-space location, required for robust estimation of cardiac-induced noise, and data acquisition was conducted for consecutive kernel positions along the slow phase encoding direction (Figure S2C).

Strategy 1 completes data acquisition at each k-space location within a short time window and minimizes low-frequency effects such as scanner drift in the data. However with strategy 1, the mitigation of spurious effects only involves k-space points within the kernel. Strategy 2 allows for a full traversal of k-space along the fast phase-encode direction before repeating data acquisition within a given kernel. Strategy 2 therefore mitigates spurious effects across a larger number of k-space points, at the expense of a longer time window to complete data acquisition at a given k-space location. The time required for the acquisition of 30 samples at each k-space location is shown in Figure S3A. Only kernels with an acquisition below 120s, highlighted in green, were considered to minimize the effect of e.g. scanner drift of head motion.

### c. Determination of the optimal mitigation strategy

#### 1. Simulating the occurrence of spurious effects

We conducted numerical simulations to identify the optimal strategy to mitigate spurious effects on the characterization of cardiac-induced noise, as follows. Each data point in Figure S1A was labelled as artefact-free or affected by spurious effects using a threshold *β* value of 5.5. We selected one data point for each label to define template signal variations across the cardiac cycle arising from cardiac-induced and spurious effects (Figure S1B). From these labels, we also computed the successive occurrence of artefact-free and spurious periods in time, given the 2D linear trajectory used to acquire the original data and the 30 repetitions acquired at each k-space location.

From the succession of artefact-free and spurious periods, we simulated a time series of cardiac-induced and spurious signal variations by appropriate sampling of the cardiac and spurious templates. At each time point, the template signals were sampled for a value of the cardiac phase taken from experimental recordings of cardiac pulsation.

In our simulations, this time series of cardiac-induced and spurious signal variations was sampled by the proposed mitigation strategies and distributed across k-space. After completion of the simulated data acquisition, the variation of the acquired data across the cardiac cycle at each k-space location was fitted with Fourier series of sinusoidal basis functions with fundamental and first harmonic.

The efficiency of each mitigation strategy in sampling cardiac-related fluctuations was measured from the percentage of k-space locations with more than 3 empty cardiac bins (Figure S3B). To assess the ability of each mitigation strategy and kernel size to distribute spurious effects, we computed the percentage of k-space locations with more than 3 spurious samples (Figure S3C). The bias induced by spurious effects on the *β* estimates was computed as the difference between the *β* estimates obtained from the simulations and the value of *β* of the original cardiac-induced template (Figure S3D). We verified the homogeneity of the bias of the *β* estimates induced by spurious effects between k-space points by computing their standard deviation (Figure S3E).

#### 2. Choosing the optimal kernel

With strategy 1, the use of small kernel sizes is equivalent to the acquisition of all 30 samples in each k-space location consecutively: cardiac fluctuations are nearly always sampled for all values of the cardiac phase, but spurious effects are focused on a subset of k-space location, leading to large bias of the *β* estimates. With increasing kernel size, the mitigation of spurious noise across k-space locations improves (Figure S3C), leading to a lower average bias *β* estimates with improved homogeneity across k-space (Figure S3D and Figure S3E). Large kernels also allow efficient sampling of cardiac-induced fluctuations (Figure S3B). Sampling strategy 2 leads to more complete mitigation of spurious noise overall, with a reduced dependence on the kernel size. Specific sets of kernel sizes lead to insufficient sampling of cardiac-induced fluctuations due to synchronization of data acquisition with cardiac pulsation.

From these results, we opted to use sampling strategy 2 with a kernel size of 30 and 2 along the fast and slow phase-encode directions as it represents a good trade-off between the mitigation of spurious effects and efficient sampling of cardiac-induced signal fluctuations.

**Figure S1:**
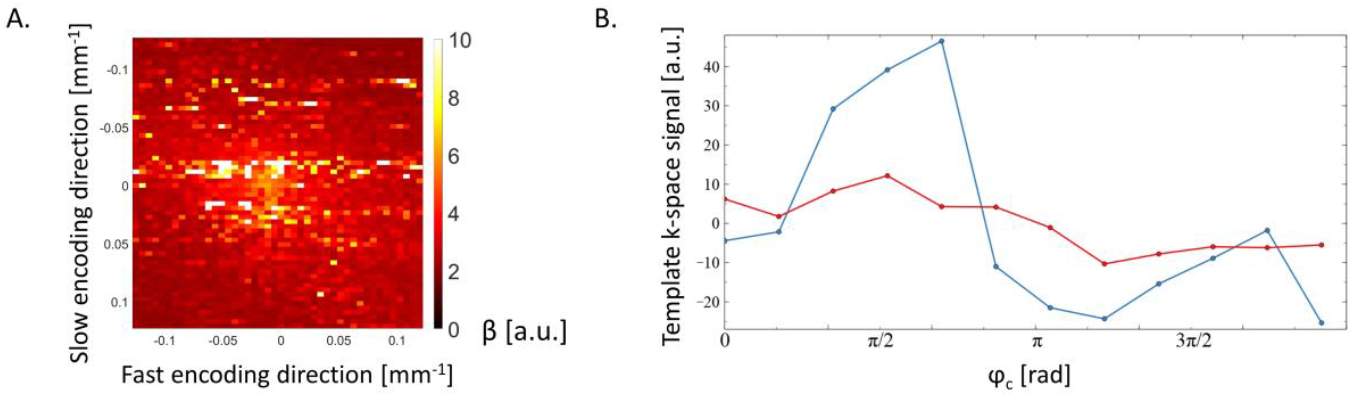
Impact of spurious effect on cardiac-induced noise modeling. (A.) Amplitude of the modeled cardiac-induced noise obtained from 30 data samples acquired consecutively at each k-space location using a 2D linear sampling. The streaks of large *β* values along the fast encoding direction arise from temporally coherent spurious noise. (B.) Template signal variations across the cardiac cycle due to spurious effects (blue) and cardiac pulsation (red).

**Figure S2:**
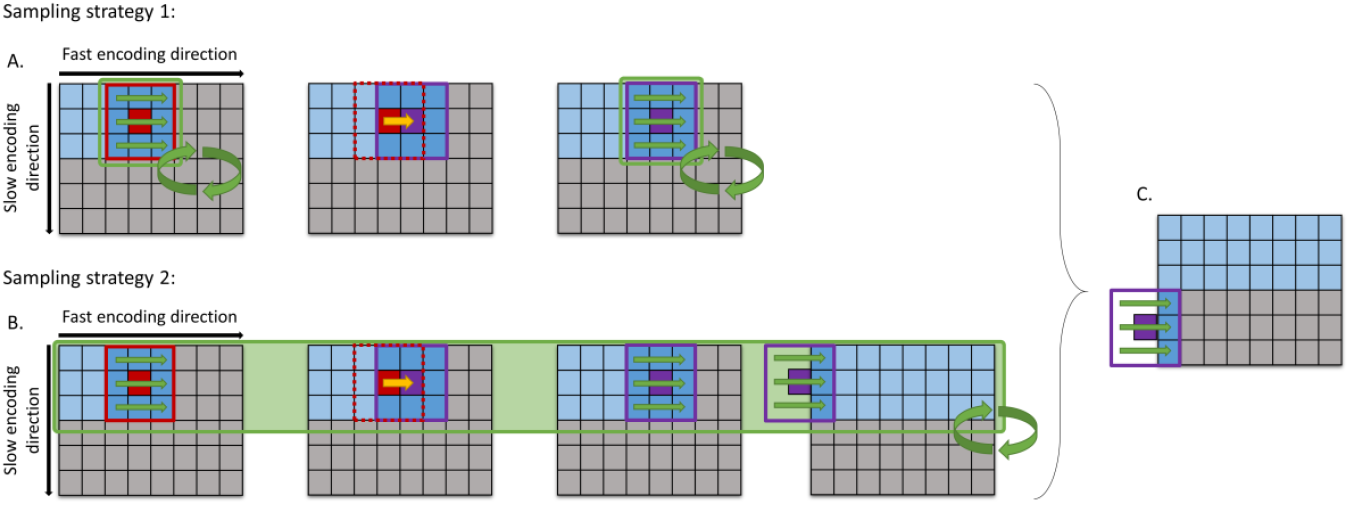
Schematic representation of the kernel displacements in k-space for the two sampling strategies. (A.) Sampling strategy 1 repeats data acquisition within the kernel several times before shifting the kernel position by one k-space index along the fast phase-encode direction. (B.) Sampling strategy 2 shifts the kernel position after each kernel acquisition and the process is repeated after completing the traversal of k-space along the fast phase-encode direction. (C.) Both strategies lead to a total of 30 samples at each k-space location, and data acquisition was conducted for consecutive kernel positions along the slow phase encoding direction. The grey squares represent k-space indexes that remain to be acquired and the blue squares represent k-space indexes that have been acquired at least once. The red and purple boxes represent two consecutive positions of the kernel. The green box represents the time span between the repetition of data acquisition at the same kernel location.

**Figure S3:**
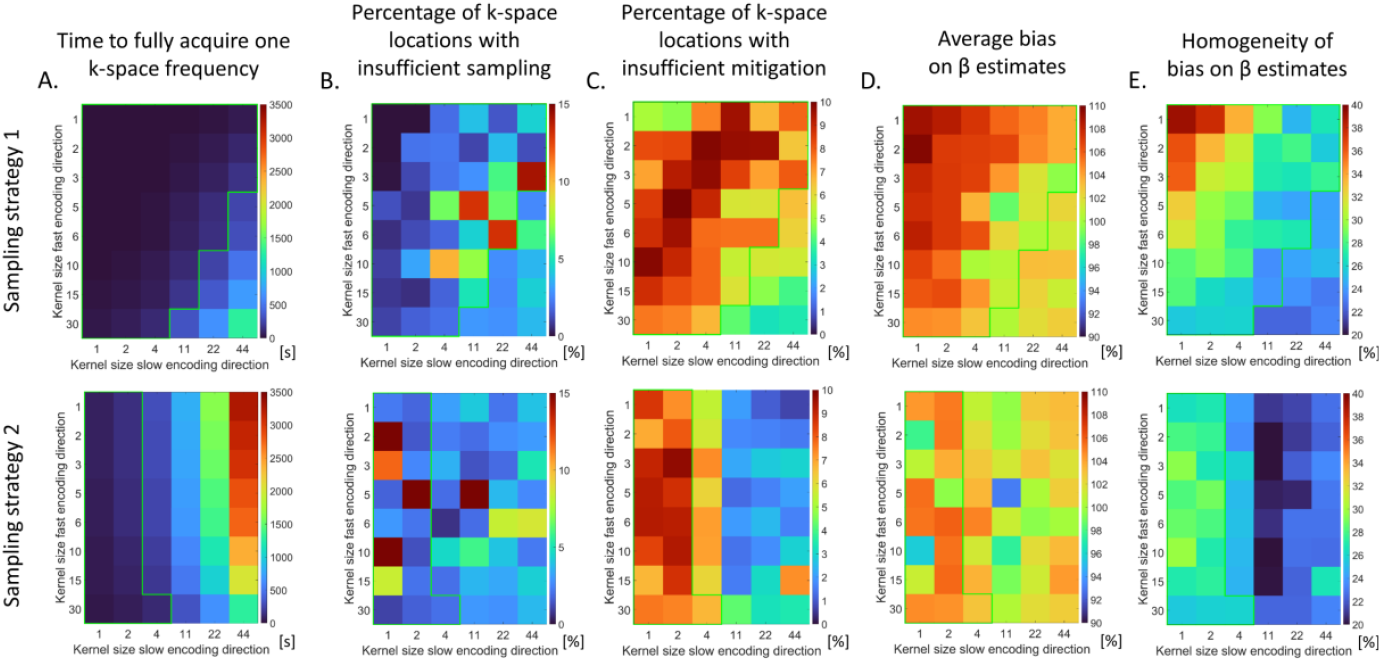
Evaluation of the sampling strategies 1 and 2. (A.) Time of acquisition of 30 samples at each k-space location. Only kernels with an acquisition time below 120s, highlighted in green, were considered. (B.) Percentage of k-space locations with insufficient sampling (i.e. more than 3 empty cardiac bins) (C.) Percentage of k-space locations with insufficient spurious effect mitigation (i.e. more than 3 spurious samples). (D.) Average bias on the *β* estimates induced by spurious effects. (E.) Homogeneity of the bias on the *β* estimates across k-space locations.

